# ResistoMap — online visualization of human gut microbiota antibiotic resistome

**DOI:** 10.1101/070714

**Authors:** Konstantin Yarygin, Boris Kovarsky, Tatyana Bibikova, Damir Melnikov, Alexander Tyakht, Dmitry Alexeev

## Abstract

**Summary:** We created ResistoMap — a Web-based interactive visualization of the presence of genetic determinants conferring resistance to antibiotics, biocides and heavy metals in human gut microbiota. ResistoMap displays the data about more than 1600 published gut metagenomes of the world populations including both healthy subjects and patients. Multiparameter display filters allow visual assessment of the associations between the meta-data and proportions of resistome. The geographic map navigation layer allows to state hypotheses regarding the global trends of antibiotic resistance and correlate the gut resistome variations with the national clinical guidelines on antibiotics application.

**Availability and implementation:** ResistoMap is publicly available at http://resistomap.datalaboratory.ru

## Introduction

Microbial drug resistance is a global problem of healthcare caused by the extensive uncontrolled use of antibiotics in medicine and agriculture: it is predicted that toward 2050 around 10 million people will die annually due to the reasons connected with drug resistance [1]. Although the main concern are resistant pathogens, the global microbial chanels of gene exchange existing between the unrelated microbial taxa allows the commensal microbes to share the resistance genes with the opportunists [2].

Human gut microbiota is being increasingly viewed at as a clinically important reservoir of drug resistance [3]. Among other human-associated communities, this one is the largest and most tightly regulating the host health. The pool of antibiotic resistance (AR) genes that is increased in abundance during antibiotic treatment becomes a “tinderbox”: the transmission of these genes to a pathogen has dire conseqeuences for both the subject and the society [4].

Semi-quantitative analysis of the functional composition of microbiota using “shotgun” metagenomics allows to assess the relative abundance of the AR genes in human microbiota (by counting the reads aligned to each gene sequence) [5, 6] and thus provide a personal prediction for the individual capacity of microbiota to contributing to the on-set of resistant pathogens. Vast volumes of metagenomes have been published that provide the opportunity to estimate the variation of resistome between the subjects, populations of the world as well as clinical cohorts. However, there is a lack of visual tools for exploratory analysis of such data; moreover, no unified actual database of gut resistome exists.

Here we are present ResistoMap, an interactive tool for comprehensive visualization of the gut resistome in the populations of the world. The displayed features include not only the relative abundance of the AR genes but also of the AR-conferring mutations as well as of the genes conferring the resistance to biocides and heavy metals. ResistoMap is a perspective tool for exploring the global landscape of gut resistome in order to identify the national traits in antibiotic intake, correlating the resistome composition with various external and generate biomedical hypotheses that can help to control the drug resistance on the global scale.

## Implementation

### Navigation

ResistoMap contains two main interactive work fields — a geographic map and a heatmap.

The heatmap displays the median relative levels of the determinants conferring resistance to each of the antibiotic groups (columns) in each of the selected cohort of subjects (rows). The values were precomputed by classifying the gut metagenomic reads from 12 publicly available studies (see Metagenomic data). The number of the metagenomes included in the cohort is displayed on the left from the heatmap below the color key.

The heatmap contains four vertical sections corresponding to different types of resistome quantification:

1. levels of the AR-conferring genes;
2. levels of the mutations in the target genes;
3. levels of the genes conferring resistance to biocides (total);
4. levels of the genes conferring resistance to heavy metal (total).

Using the drop-down lists at the top of the screen, a user can choose the antibiotic group of interest, the study(-ies) and/or the country(-ies) to be displayed at the heatmap. For filtering the cohort, the user can use “Pool by” checkboxes that allow to stratify the cohorts by the country of origin, gender, age and diagnosis (where applicable).

### Metagenomic data

The datasets for the analysis of resistome included 1638 gut metagenomes from the individuals from 12 studies covering 15 countries [7, 8, 9, 10, 11, 12, 13, 14, 15, 16, 17, 18]. For each metagenome, additional factor including the country of origin, gender, age and clinical status were considered (where available).

Using ResistoMap, one can estimate the global variation of the resistance potential to different groups of antibiotics and explore the associations between specific drugs and the clinical factors and other meta-data. For instance, the Danish gut metagenomes tend to demonstrate the lowest resistome among the European groups, whereas the French samples have the highest levels, particularly, of the fluoroquinolones. It is in agreement with the fact that France has the highest total antibiotic use within Western Europe [19], while Denmark and Germany are countries where the use of antimicrobial drugs is moderate in both health-care and agriculture [20]. On the other hand, Chinese and Russian populations appear to have increased levels of resistome — likely to more relaxed respective regulations, frequent prescription of wide-spectrum antibiotics as well as over-the-counter availability of the antibiotics without the prescription [21, 22]. The lowest levels of microbiota resistome are observed in Amerindians from Venezuela who have no documented contacts with Western people. Strikingly, 1-month old childrens from Singapore have the highest resistome among all groups — this fact might be associated with the potential antibiotic treatment of the newborn.

Some of the results obtained using ResistoMap are in agreement with the earlier analyses performed for smaller collection of datasets and mirror certain aspects of national specifics of antibiotic use. Moreover, the analysis reveals certain novel trends that await further interpretation from the clinical perspective. We suggest that the exploratory analysis of global gut resistome using ResistoMap will gain new insights approaching the development of efficient antibiotic stewardship and agricultural use of antimicrobial substances.

## Methods

### Data processing

The reference nucleotide sequences of the resistance-conferring genes were taken from the CARD database v.1.0.5 [23]. The metagenomic reads were mapped to the reference set using Bowtie2 [24] using the parameters *-k 1* (best hit) (for Illumina reads) and Bowtie [25] using the parameters *-v 3 -k 1* (for SOLiD reads [10]). The relative abundance for each gene was calculated by normalizing the gene coverage by gene length and the total number of the reads in the metagenome (*gene abundance* = *count/*(*L * N)*). The resistance potential to each group of antibiotics (according to CARD) was calculated as the sum of the abundance of all the genes conferring resistance to these drugs.

For the analysis of the potential resistance caused by mutations in the target genes, we used SAP (single amino-acid polymorphism) table [26]. For the genes *folP, gyrA, gyrB, parC, parE and rpoB*, all possible nucleotide and amino-acid sequences were taken from PATRIC database [27]. For every gene, a multiple alignment was performed by applying MUSCLE v3.8.31 [28] to all of its amino-acid sequences to identify the positions that can potentially confer drug resistance. The metagenomic reads were mapped to the gene sequences using Bowtie2 software using the parameters *-k 1* (best hit). The relative abundance for each gene was calculated as described above. The relative abundance of the potentially resistant mutants among the microbial populations were calculated as the gene relative abundance multiplied by the share of the SAPs potentially conferring resistance: *mutant gene abundance* = *gene abundance * alt/*(*alt* + *ref*). Then the resistance potential to each group of antibiotics was calculated by summing.

The reference amino-acid sequences of genes providing resistance to heavy metals and biocides were taken from BacMet database [29]. The metagenomic reads were mapped to the reference set using DIAMOND [30] with the parameters *-k 1* (best hit). Gene abundance were calculated by normalizing the gene coverage by gene length and the total number of the reads in the metagenome (*gene abundance* = *count/*(*L*N)*). The relative abundance for each gene and the total resistance potential to each of the metals or biocides group were calculated as described above.

